# Attenuation of endothelial glycocalyx shedding and endocan modulation by Sulodexide in murine models of anaphylaxis

**DOI:** 10.1101/2025.10.16.682838

**Authors:** Lucía Palacio-García, Sergio Fernández-Bravo, Irene de María-Camacho, Irene San Sebastián-Jaraba, María José Fernández-Gómez, Óscar López-Pérez, Sandra Sanz-Andrea, Carlos Pastor-Vargas, Emilio Nuñez-Borque, José Julio Laguna, Marcela Valverde, Luis Miguel Blanco-Colio, Vanesa Esteban, Nerea Méndez-Barbero

## Abstract

**Background:** Anaphylaxis is an acute life-threatening reaction. Research into the vascular endothelium and its components may improve disease management and patient outcomes.

**Objective:** We investigated the endothelial glycocalyx (eGCX) and its pathophysiological role in murine anaphylaxis, aiming to identify novel diagnostic and therapeutic targets.

**Methods:** Active systemic anaphylaxis (ASA) and passive systemic anaphylaxis (IgE-PSA and IgG1-PSA) models were evaluated in mice. Sulodexide (Sdx) was administered as a prophylactic treatment. eGCX structure and N-acetylglucosamine residues in mouse aortic tissue were analyzed by electron microscopy and wheat germ agglutinin (WGA) staining. Endocan (ESM-1) levels in mouse aorta and plasma were determined by immunofluorescence and ELISA. Human sera from beta-lactam-induced anaphylaxis and endothelial cell (EC) secretome samples were also analyzed.

**Results:** ASA, IgE-PSA, and IgG1-PSA models showed reduced eGCX surface area and thickness. N-acetylglucosamine and ESM-1 levels decreased in aortic tissue but increased in plasma, indicating glycocalyx shedding. Consistently, ESM-1 secretion was enhanced in ECs exposed to acute anaphylactic sera. ESM-1 and hyaluronic acid levels differed significantly between anaphylactic patients and non-allergic controls. Sdx reduced reaction severity in ASA and IgE-PSA, increased survival in ASA, and prevented eGCX disruption and ESM-1 release.

**Conclusions:** eGCX shedding, particularly of ESM-1, acts as a key mediator in murine anaphylaxis. Sdx prophylaxis protects against severe reactions and improves survival.

**Clinical Implication:** Therapies based on glycosaminoglycans and proteoglycans may mitigate anaphylaxis severity, and monitoring eGCX dynamics could aid diagnosis.

**Capsule summary:** Endothelial glycocalyx shedding contributes to anaphylaxis pathophysiology; targeting its preservation and measuring HA and ESM1 may offer novel diagnostic and therapeutic strategies for clinicians.

## INTRODUCTION

Anaphylaxis is a life-threatening systemic hypersensitivity reaction caused by different elicitors.(1) The increased vascular permeability and extravasation of fluids that characterise severe reactions underscore the central role played by the endothelium in anaphylaxis pathophysiology. (2,3) This large organ participates in numerous vital functions, including coagulation and fibrinolysis modulation, the regulation of immune-cell transport, control of cellular metabolism, and the release of angiocrine mediators.(4) During anaphylaxis, the endothelial layer is affected by multiple molecular agents triggered by immune cell activation and homeostasis imbalance.(5) This leads to endothelial-barrier breakdown within seconds, causing extravasation of vascular contents into the surrounding tissues.(6) Consequently, intravenous (i.v) fluid administration is recommended for acute management.(7,8)

The endothelial cell (EC) surface is covered by a dynamic, shifting structure primarily composed of glycoproteins and proteoglycans, including glycosaminoglycans (GAG) side chains, collectively termed as the endothelial glycocalyx (eGCX).(9,10) This malleable sugar-based structure acts as a signalling platform and is essential for communication between tissues and the bloodstream. Its plasticity makes it highly prone to damage, which leaves the endothelial membrane unprotected and triggers endothelial dysfunction associated with pro-inflammatory conditions such as diabetes, hypertension, sepsis, or atherosclerosis (11–13).

The eGCX controls leukocyte adhesion and transmigration through endothelial activation of surface receptors such as vascular adhesion molecule 1 (VCAM-1), among others (14). Damage to the eGCX results in pathological manifestations including capillary leak, augmented migration of pro-inflammatory cells, and increased endothelial nitric oxide (NO) synthase activity.(15,16) Therefore, preserving the structural integrity of the eGCX is crucial to maintain endothelial stability by limiting the passage of certain molecules to surrounding tissues and preserving homeostasis.(17)

Strategies aimed to prevent eGCX degradation hold therapeutic potential.(18,19) Sulodexide (Sdx), a precursor of GAG synthesis mainly composed of heparin and dermatan sulfate, modulates eGCX remodelling (20), thus emerging a promising approach. With established anti-inflammatory, anti-thrombotic, and profibrinolytic activity, it is primarily used in clinical settings to treat peripheral arterial diseases and chronic venous disorders as an alternative to oral heparin or anticoagulant.(21–23) Furthermore, Sdx has demonstrated efficacy in eGCX remodelling in sepsis.(19)

eGCX shedding induces local structural alterations leading to the release of its components into the bloodstream, modulating signalling pathways in an endocrine manner. Accordingly, circulating eGCX components are potential biomarkers for diagnosing endothelial dysfunction. (9)(24) Among the specific constituents of the eGCX, endocan, or endothelial cell specific molecule-1 (ESM-1), is a soluble dermatan sulfate proteoglycan secreted by vascular ECs. Several reports have demonstrated that *esm-1* mRNA expression is regulated by various molecules including tumour necrosis factor-α (TNF-α), interleukin-1β (IL-1β), interleukin-8 (IL-8), E-selectin, transforming growth factor-β1 (TGF-β1), lipopolysaccharide (LPS), nuclear factor-κB (NF-κB), and vascular endothelial growth factor (VEGF).(24–28) ESM-1 plays a crucial role in several diseases by enhancing vascular permeability, leukocyte adhesion, and production of pro-inflammatory cytokines.(29–31) ESM-1, along with other eGCX components such as syndecan-1, heparan sulfate, and hyaluronic acid (HA), serves as a serological biomarker of inflammatory conditions related with eGCX injury. Increased degradation of the endothelial surface into the bloodstream correlates with disease severity in various pathological settings.(32) Elevated blood levels of ESM-1 have been observed in autoimmune diseases, acute inflammatory conditions such as sepsis, and cardiovascular diseases, corroborating its status as a novel biomarker(33–37) as well as a therapeutic target.(38)

We aimed to expand the understanding of eGCX integrity in anaphylaxis, a crucial factor in regulating associated endothelial dysfunction, and explore eGCX protection as a potential therapeutic strategy. To achieve this, we analysed the rapid shedding of eGCX and evaluated ESM-1 as a diagnostic tool. In addition, we explored the prophylactic function of Sdx in anaphylactic mice models.

## METHODS

### Animal experimental design

All procedures using animals were conducted in accordance with European Union Directive 2010/63/EU and Recommendation 2007/526/EC for the care and experimental use of animals. The experimental protocols used in the study (reference code PROEX: 074.4/21) were approved by the local Ethics Committee and the Comunidad de Madrid prior to use. We developed mouse models of active systemic anaphylaxis (ASA) by active immunization and passive systemic anaphylaxis (PSA) induced by transfer of antigen-specific IgE- or IgG and subsequent challenge into naive mice as previously described.(39). 8-to-12-week-old animals of the C57BL/6 strain (Charles River Laboratories) were used. Expanded methodology is described in the **Supplementary Materials**. Challenged, non-sensitised mice were used as controls. The onset of anaphylactic reaction was evaluated by monitoring rectal body temperature with a digital thermometer (VWR l60-200) before challenge and at 5-minute (min) intervals following the reaction. Two hours before the challenge, 40 mg/kg of Sdx (AlfaSigma) was administered by oral gavage in select groups of mice.(19) Surviving mice were euthanised with CO_2_ and, immediately, a small blood volume was collected for hematocrit determination using a microhematocrit system (HemataStat II, EKF Diagnostics). Plasma and tissue samples were collected and processed for molecular analysis.

### Electron microscopy

eGCX degradation was assessed in aortas of mice and *in vitro* human aortic endothelial cells (HAEC) by transmission electron microscopy (TEM) as indicated **in the Supplementary Materials**. *ImageJ* software was used to quantify the eGCX. Both the thickness of the eGCX and the total area covered by the glycocalyx were quantified in different areas of ECs in the slide. To measure thickness, 6 sections of each sample were evaluated in a blinded manner. In turn, 6-7 measurements per image were quantified to obtain an average value per mouse.

### Tissue staining and immunofluorescence

Aortas obtained from mice were embedded in paraffin, cut into sections (4-µm thick), deparaffinised, and rehydrated before antigenic unmasking was performed using a pressure cooker and citrate buffer (pH 6). To prevent non-specific binding, blocking was done using phosphate buffer with 5% bovine serum albumin (BSA), 10% goat serum, and mouse Fc Block (1:100; BD Pharmingen). Anti-ESM-1 and anti-CD31 primary antibodies were used at a dilution of 1:50 (Abcam). Donkey anti-rabbit Alexa Fluor 488 (Invitrogen) was used for detection. Staining with fluorescent Alexa Fluor 555 wheat-germ agglutinin (WGA; Invitrogen) was carried out at a dilution of 1:2000. Computer-assisted image analysis was performed with *ImageJ* software (version 1.0, Windows). The same threshold setting for area measurement was used for all images. Samples from each animal were examined in a blinded manner. Results were expressed as % positive area vs medial area for ESM-1 staining. The intensity profile for WGA-stained images was generated using *ZENN 3.1 blue edition* software (Carl Zeiss, Canada). WGA intensity (Y-axis, au) was mapped against distance (X-axis) along a spatial line, drawn from the lumen to the medial layer of the artery (50-µm deep in histogram and 100 µm in quantification). The limit between the intimal and medial layer was defined by CD31 staining (endothelium). Measurements were performed in 3-4 independent ECs per mouse.

### RNA extraction and RT-qPCR

RNA from aortas was extracted using Tri-Reagent (Molecular Research Center). Two micrograms of total RNA was reverse-transcribed according to the high-capacity cDNA reverse transcription kit (Applied Biosystems) protocol. Samples were then stored at −20°C. Quantification of *vcam1*, and *18s* rRNA was performed with TaqMan gene-expression assays from Applied Biosystems. Quantification of *esm-1, synd-1, synd-4, acan, gpc1, hs2st1, ndst2, selp* and *18s rRNA* expression was performed with the SYBR Green detection method. All primers were purchased from Eurofins Genomics according to the sequences described in the **Supplementary Materials**. Double delta Ct analysis was performed in all datasets, and heatmaps were generated using the R programme with a specific guideline given by "Bioinformatics for all" (https://bioinfo4all.wordpress.com).

### Anaphylactic serum-sample recruitment and collection

Paired samples were obtained from 22 patients under 2 conditions: during the acute phase (anaphylaxis induced by beta-lactams) and at a baseline phase at least 14 days after the reaction. Following the criteria established by the National Institute of Allergy and Infectious Diseases and the Food Allergy and Anaphylaxis Network,(40) 13 patients experienced grade-2 reactions, and 9 experienced grade-3 reactions according to the severity grading.(41) 21 control samples from non-allergic subjects were also used. For serum collection, the blood samples were centrifuged at 1200 g for 10 min at 4°C. Serum aliquots were stored at -80°C until use. The diagnosis, selection, and blood collection of all individuals was carried out in the Allergy and Emergency Departments of Fundación Jiménez Díaz University Hospital and Cruz Roja Central Hospital. Sample collection was performed as quickly as possible at all times, prioritising patients’ lives and applying all appropriate protocols and drug administration when necessary. The project was conducted in accordance with the ethical protocols approved by the Clinical Trials Committee (CEIC-FJD, PIC057-19 and PIC166-22_FJD) and samples were collected after signed informed consent was obtained from donors.

### *In vitro* human endothelial/sera system

Human dermal microvascular endothelial cells (HMVEC-d) were seeded on P6 plates and FBS depleted 18 h before experimentation. Subsequently, cells were incubated with paired patients’ sera (acute and baseline phases independently) at a 1:2 concentration with EGM medium without supplements, to mimic an *in vitro* microenvironment that allows the release and interchange of mediators between ECs and patient sera. The former, including the cellular secretome, were collected after 2 h of incubation, aliquoted, and stored at -20°C to properly measure ESM-1 protein level

### *In vitro* anaphylactic mediators TEM assay

Human dermal microvascular endothelial cells (HAEC) were seeded in poly-L-lysine-coated glass coverslips and FBS depleted 18 h before experimentation. Then, cells were incubated with anaphylactic mediators (histamine (1µm), PAF (10µm) and thombin (1Ud) for 2h. The secretome was collected and the culture was fixed for eGCX TEM visualization as indicated in **Supplementary Materials.**

### ELISAs

Sandwich ELISAs for human ESM-1 (Elabscience) and HA (Cusabio) were used to quantify the abundance of those proteins in sera from anaphylactic patients. A sandwich ELISA for mice ESM-1 (Cusabio) was employed to measure the ESM-1 protein levels in plasma of mice experiencing anaphylaxis. In both cases, colorimetric detection was performed with the Infinite F200 system (TECAN) at a wavelength of 450 nm, following the manufacturer’s instructions.

### Statistical analysis

Data are expressed as mean ± the standard error of the mean (SEM). For mice models, statistical significance between two groups was assessed using Student’s t-test. For multiple group comparisons, the one-way or two-way ANOVA test followed by Tukey’s analysis (parametric) was used. Correlation between variables was evaluated using Pearson’s correlation coefficient.

For patients’ samples, normality was evaluated following the Shapiro-Wilk test and depending on the distribution, parametric (Student’s t-test or ANOVA) or non-parametric (Mann-Whitney or Krustal-Wallis) tests were applied as appropriate.

Statistical significance was set at *P* values <0.05. Data analysis was performed using *GraphPad Prism 8.0* software (San Diego, CA).

## RESULTS

### ASA induces endothelial glycocalyx disruption and aortic glycoprotein erosion

To evaluate possible eGCX alterations in anaphylaxis, we examined the aortic endothelium of an *in vivo* ASA mouse model (**Figure 1A**). Continuous body-temperature monitoring revealed a significant reduction (approximately 3°C) within 5 to 10 min of challenge (**Figure 1B**). Next, we conducted the comprehensive TEM analysis of the aortic eGCX architecture. Representative images of the eGCX in control mice revealed dense hair-like structures uniformly covering the endothelial surface, whereas these structures were markedly reduced or irregular in the aortas of ASA mice (**Figure 1C**). Quantitatively, we observed a 50% reduction in eGCX surface area and a 60% decrease in thickness in the aortic eGCX of ASA mice compared with controls, indicating a loss of eGCX integrity in anaphylaxis (**Figure 1D**). In addition, WGA staining revealed a reduction in glycoprotein content throughout the aortic tissue (**Figure 1E and Supplementary Figure 1A**). WGA intensity profile demonstrated a significant decrease in the endothelium and medial layer of the aortas of ASA mice, (**Supplementary Figure 1A and B**). In addition, since WGA intensity did not correlate with severity (**Supplementary Figure 1C),** hematocrit levels negatively correlate with WGA intensity and positively with the drop of body temperature in ASA mice (**Figure 1F and Supplementary Figure 1D).**

**Figure 1.**
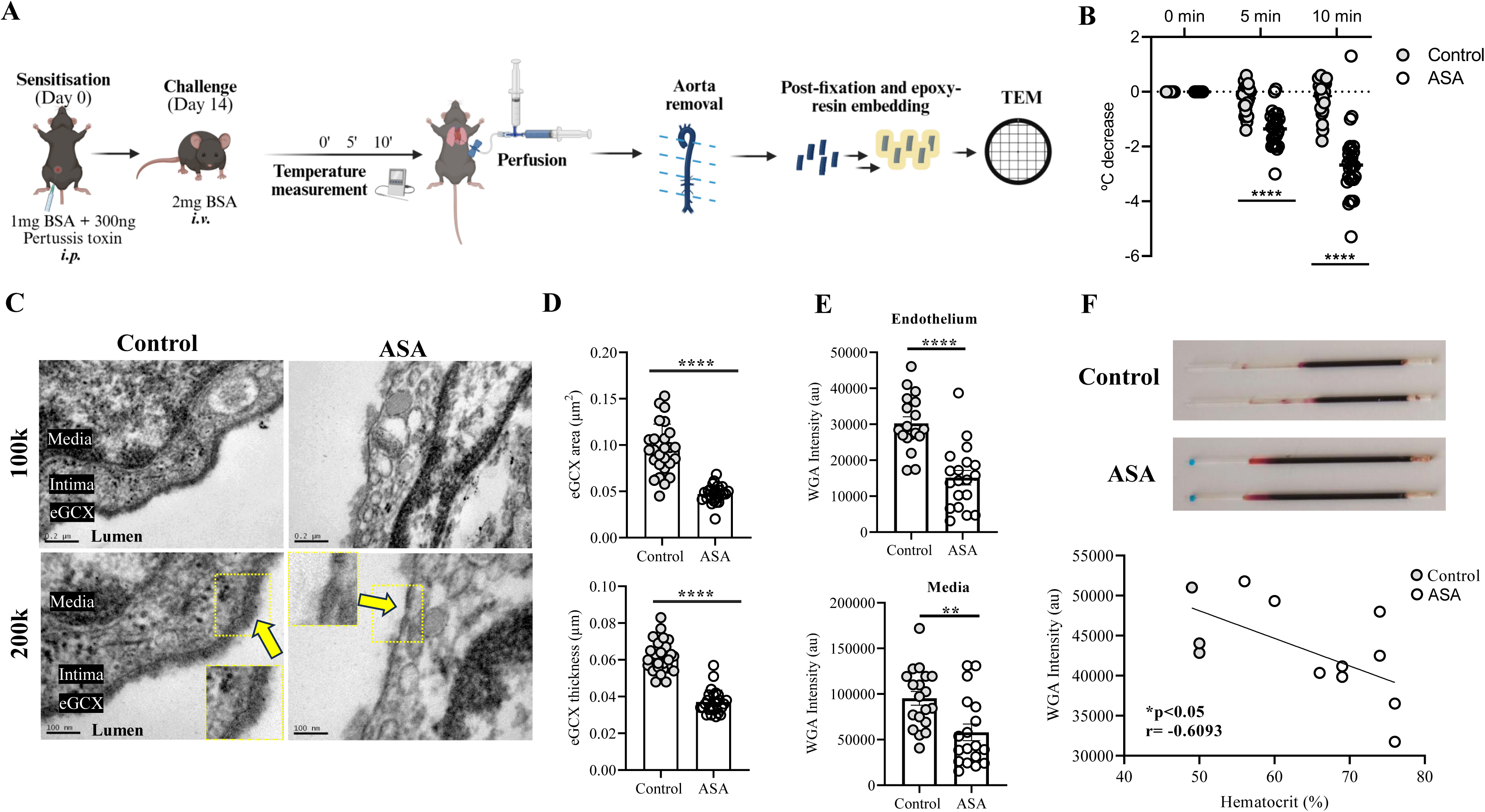
Anaphylaxis triggers endothelial glycocalyx shedding in a mouse model of active systemic anaphylaxis. **A.** Schematic representation of the ASA model and TEM procedure. **B.** Rectal temperature recording over time in control (n=26) and ASA (n=27) mice. **C.** Representative TEM images of aortas from control mice and ASA mice at different magnifications. Yellow arrows point to eGCX. **D.** Quantification of eGCX surface area (µm^2^) and thickness (µm) in control (n=4) and ASA (n=4) mice aortas. **E**. Quantification of WGA average intensity throughout the endothelium and medial layer. Three profiles per animal from control (n=6) and ASA (n=6) mice. **F**. Representative hematocrit capillaries images and correlation between hematocrit percentage and endothelial WGA intensity staining. Correlation was performed in Control (n=3) and ASA (n=9) mice.BSA: bovine serum albumin, TEM: transmission electron microscopy, ASA: active systemic anaphylaxis, eGCX: endothelial glycocalyx, WGA: wheat germ agglutinin. (\*\**P*<0.01, \*\*\**P*<0.001, \*\*\*\**P*<0.0001)

### Sulodexide prophylaxis prevents mice survival, drop of body temperature and endothelial glycocalyx disruption

To investigate the effect of eGCX shedding prevention in anaphylaxis, active immunized mice were prophylactically treated with Sdx (**Figure 2A**). While ASA mice died within 20 min of challenge, 55% of the Sdx-treated ASA animals survived, indicating a marked improvement in survival of this group (**Figure 2B**). Similarly, ASA challenge induced a reduction of 2 degrees in body temperature at 5 min and 4 degrees at 10 min post-challenge, while Sdx administration significantly attenuated the drop in Sdx-treated ASA mice (**Figure 2C**). In addition, aortic exploration in prophylactically treated mice showed the prevention exerted by Sdx in eGCX area shedding and endothelial WGA intensity, (**Figure 2D-F and Supplementary Figure 2**). Hematocrit levels were not modified by Sdx (**Figure 2G**).

**Figure 2.**
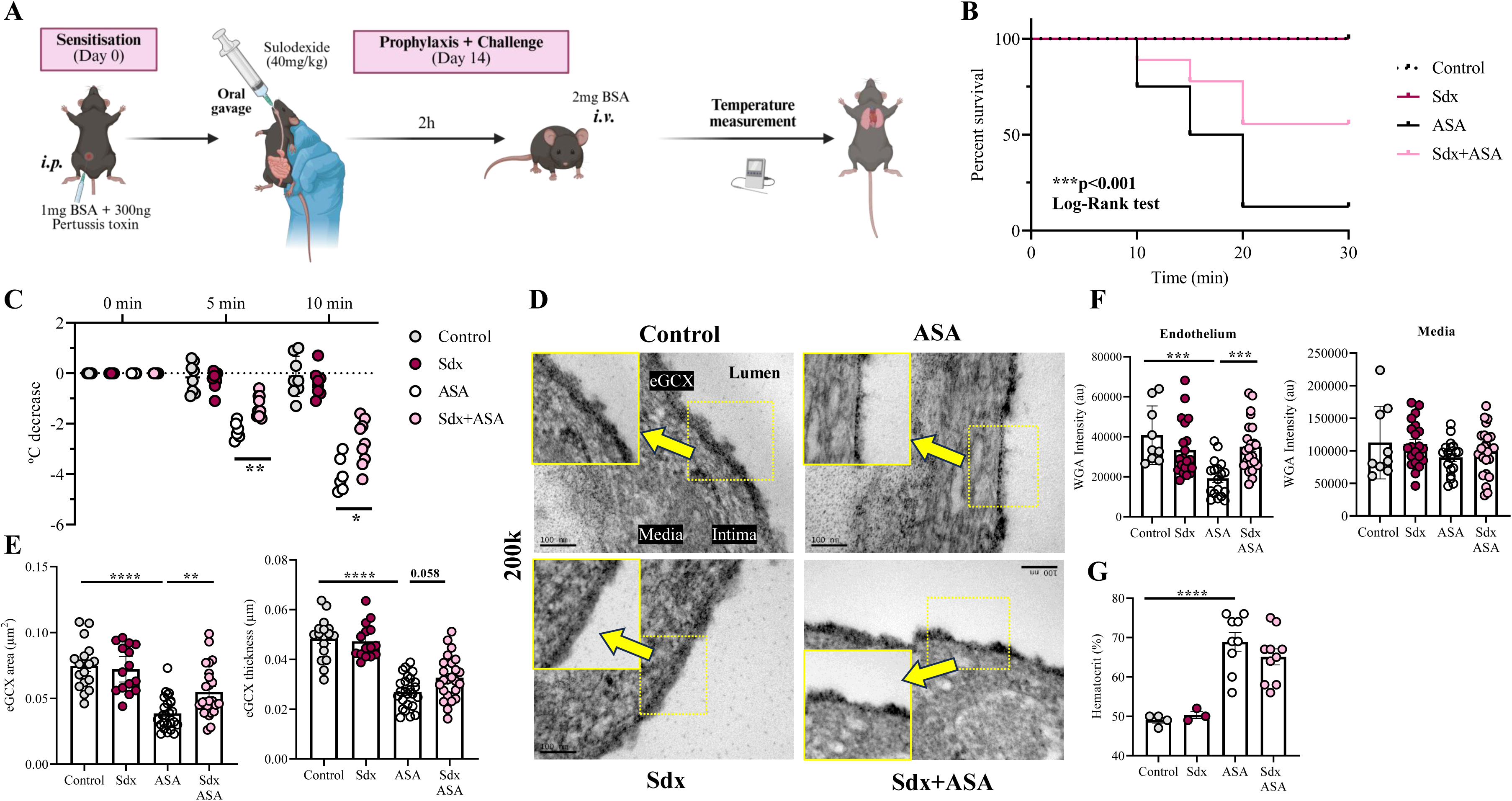
Prophylactic treatment with sulodexide improves survival, attenuates anaphylaxis severity and prevents eGCX shedding in aortas of active systemic anaphylaxis mice. **A.** Schematic representation of the ASA model including Sdx prophylaxis. **B**. Survival curve. Control (n=8), Sdx (n=8), ASA (n=6) and Sdx+ASA (n=9) mice. **C.** Rectal temperature recordings after challenge represented as degree of decrease. Control (n=8), Sdx (n=8), ASA (n=6) ASA+Sdx (n=9). **D**. Representative TEM images of aortas from control, Sdx, ASA and Sdx+ASA mice at 200k magnification. Yellow arrows point to eGCX. **E**. Quantification of eGCX surface area (µm^2^) and thickness (µm) in control (n=4), Sdx (n=3), ASA (n=5) and Sdx+ASA (n=5) mice aortas. **F**. Quantification of WGA average intensity throughout the endothelium and medial layer. Three profiles per animal from control (n=3), Sdx (n=7), ASA (n=6) and Sdx+ASA (n=8) mice. **G**. Hematocrit percentage in control (n=4), Sdx (n=3), ASA (n=9) and Sdx+ASA (n=10) mice. BSA: bovine serum albumin, ASA: active systemic anaphylaxis, IP: intraperitoneal, IV: intravenous, eGCX: endothelial glycocalyx, WGA: wheat germ agglutinin, Sdx: sulodexide. (*=*P*<0.05, **=*P*<0.01, \*\*\**P*<0.001, \*\*\*\**P*<0.0001).

### Endothelial cell-specific molecule-1 plasma levels correlate with ASA severity

Next, we evaluated the transcriptomic profile of specific endothelial activation and eGCX related genes in total aortic homogenates. The analysis exhibited a quick and significantly increased mRNA expression of *esm-1, selp* and *vcam1* compared to aortas from control mice (**Figure 3A and Supplementary Figure 3A-B)**. However, the expression of *acan*, *sydn-1, sydn-4*, *ndst2*, *gpc1*, and *hs2st* remained unchanged at this time (**Supplementary Figure 3**). As our main research focus was the eGCX, we characterised the ESM-1 protein. Staining revealed pronounced shedding throughout the aortic wall in ASA mice, (positive staining of 49% in controls vs 17% in ASA mice) (**Figure 3B-C**) that was regulated by Sdx treatment, indicating a prevention of shedding in aortas of Sdx-treated ASA mice (**Figure 3C and Supplementary Figure 3C**). Accordingly, ESM-1 plasma levels from ASA mice were increased after 10 min of challenge, and significantly prevented in Sdx-treated ASA mice (**Figure 3D)**. In addition, ESM-1 plasma levels showed a positive correlation with the degree of body temperature reduction, indicating an association between ESM-1 levels and temperature change (**Figure 3E**).

**Figure 3.**
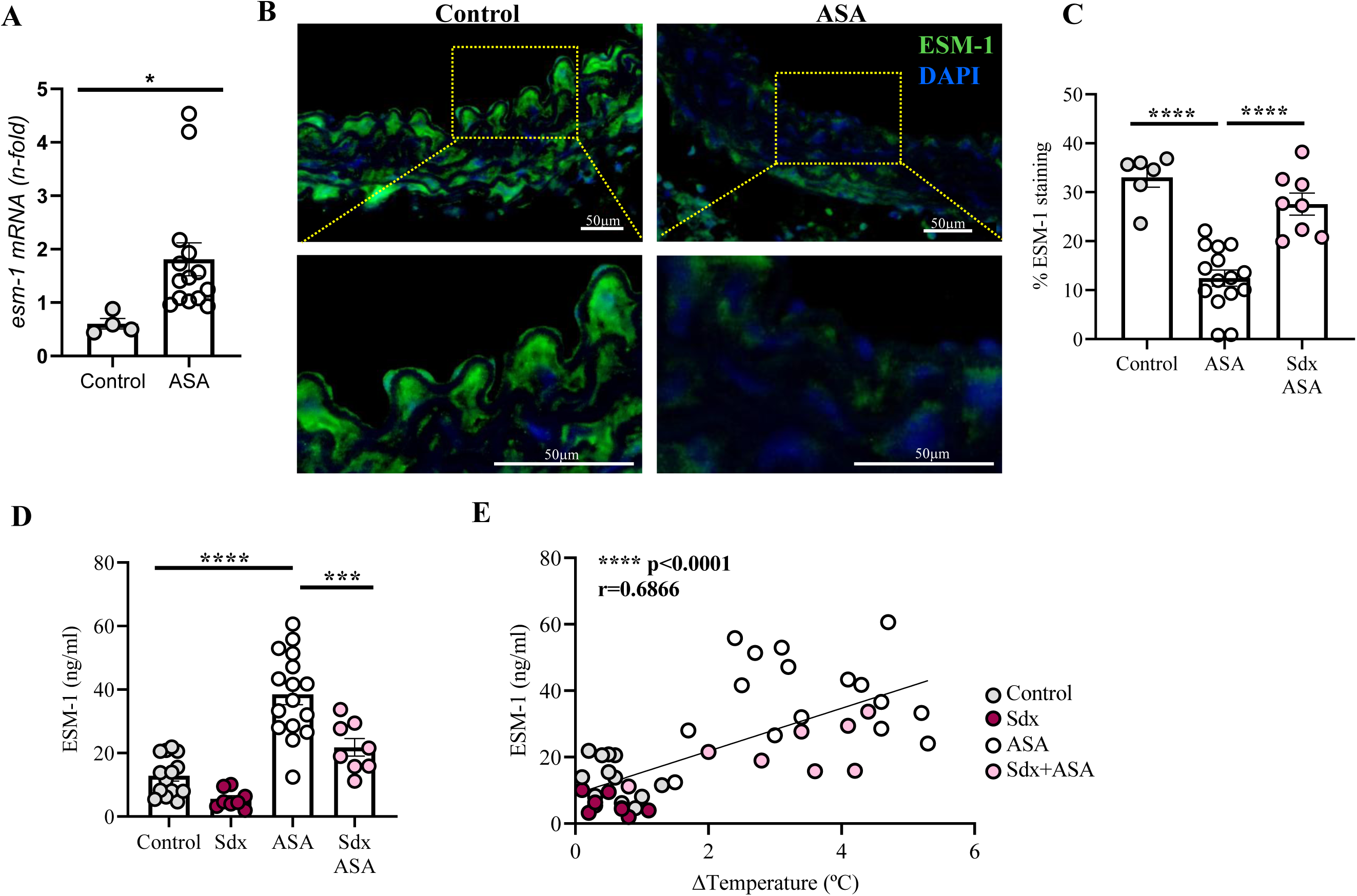
Endothelial cell-specific molecule-1 plasma levels correlate with ASA severity and it is prevented by Sulodexide. **A**. *esm-1* RNA expression in aortas from control (n=4) and ASA (n=14) mice. **B**. Representative images of ESM-1 (green) and DAPI (blue) staining in aortic samples of control and ASA mice. **C**. Percentage of ESM-1 positive staining per area in the entire aortic layer (endothelial and media layer) from control (n=6), ASA (n=15) and Sdx+ASA (n=8) mice. **D**. Plasma levels of ESM-1 in control (n=14), Sdx (n=8), ASA (n=16) and Sdx+ASA (n=8) mice. **E**. Correlation between temperature decrease in degrees and ESM-1 plasma levels in control (n=14), Sdx (n=8), ASA (n=16) and Sdx+ASA (n=8) mice. ESM-1: endothelial specific molecule 1, ASA: active systemic anaphylaxis, Sdx: sulodexide. (\**P*<0.05, \*\*\**P*<0.001, \*\*\*\**P*<0.0001).

### IgE- and IgG1- passive systemic anaphylaxis dysregulates the eGCX and induces aortic glycoprotein erosion

To enlarge the mechanistic understanding of eGCX shedding, IgE-PSA and IgG1-PSA mice models were used. Continuous body-temperature monitoring revealed a significant reduction along the time in both groups of passively sensitized mice. Interestingly, Sdx-treated IgE-PSA mice presented prevention of drop in body temperature, but no prophylactic effect was observed in Sdx-treated IgG1 PSA animals (**Figure 4A-B**).

**Figure 4.**
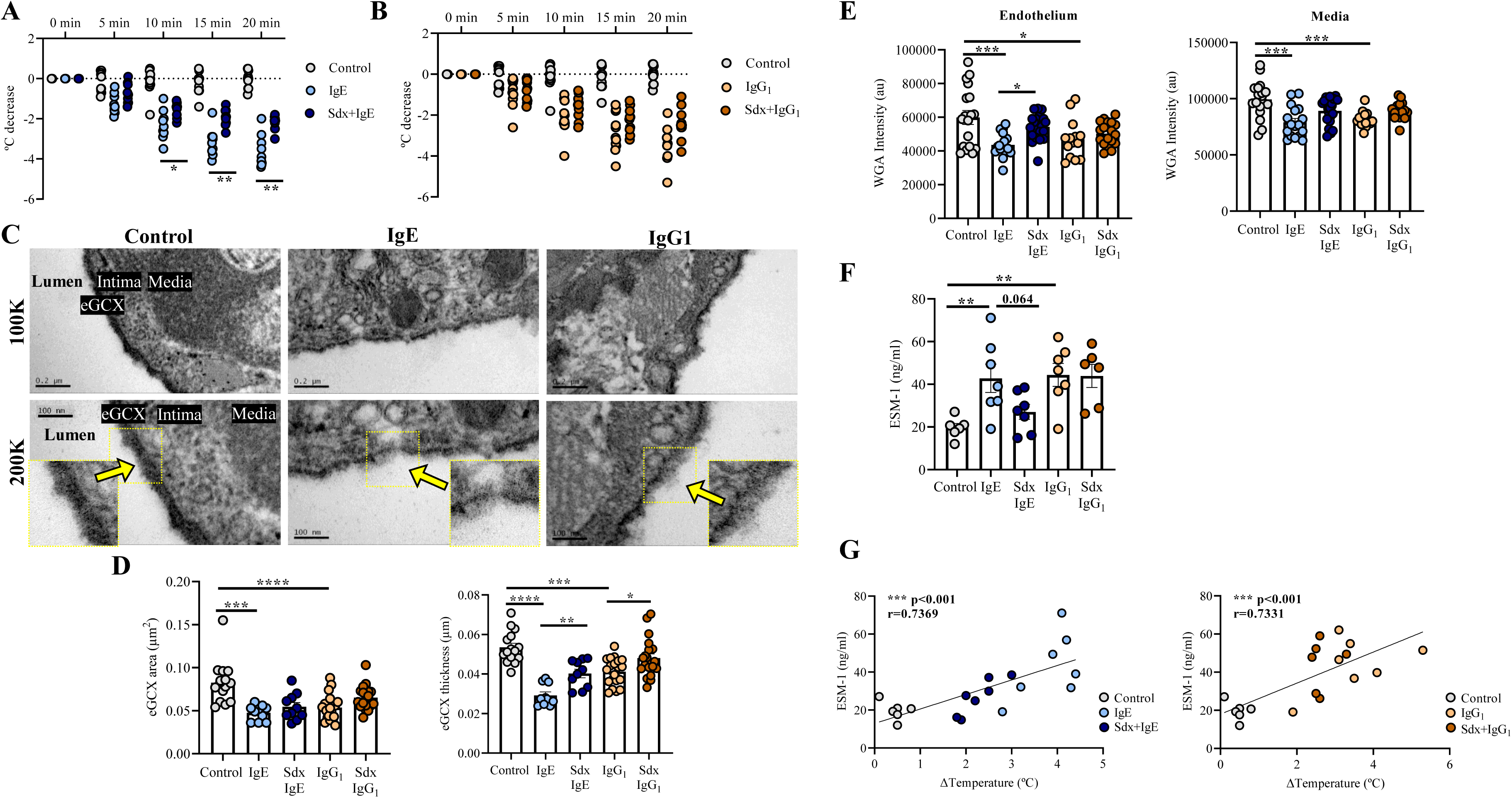
Endothelial glycocalyx dysregulation in IgE- and IgG1-dependent passive systemic anaphylaxis. **A.** Rectal temperature recordings after challenge represented as degree of decrease in IgE-dependent PSA model. Control (n=10), IgE (n=9) Sdx+IgE (n=9). **B.** Rectal temperature recordings after challenge represented as degree of decrease in IgG1-dependent PSA model. Control (n=10), IgG1 (n=11) Sdx+IgG1 (n=10). **C**. Representative TEM images of aortas from control, Sdx, IgE and IgG1 mice at different magnifications. Yellow arrows point to eGCX. **D**. Quantification of eGCX surface area (µm^2^) and thickness (µm) in control (n=3), IgE (n=2), Sdx+IgE (n=2), IgG1 (n=4) and Sdx+IgG1 (n=4) mice aortas. **E**. Quantification of WGA average intensity throughout the endothelium and medial layers. Three profiles per animal from control (n=6), IgE (n=6), Sdx+IgE (n=7), IgG1 (n=5) and Sdx+IgG1 (n=6) mice. **F.** Plasma levels of ESM-1 in control (n=6), IgE (n=7), Sdx+IgE (n=7), IgG1 (n=7) and Sdx+IgG1 (n=6) mice. **G**. Correlation between temperature decrease in degrees and ESM-1 plasma levels in IgE- and IgG1-dependent PSA models. Control (n=6), IgE (n=7), Sdx+IgE (n=7), IgG1 (n=7) and Sdx+IgG1 (n=6) mice. PSA: passive systemic anaphylaxis, eGCX: endothelial glycocalyx, TEM: transmission electronic microscopy, WGA: wheat germ agglutinin, Sdx: sulodexide, ESM-1: endothelial specific molecule 1. (*=*P*<0.05, **=*P*<0.01, \*\*\**P*<0.001, \*\*\*\**P*<0.0001).

TEM analysis showed a marked reduction in eGCX surface area and thickness in both IgE-PSA and IgG1-PSA. Specifically, IgE-PSA mice exhibited structural eGCX alterations with reduced thickness and density, yet the eGCX remains continuous, unlike in the ASA mice, where it is partially absent. In contrast, IgG1-PSA mice maintain a more preserved eGCX structure, with slightly reduced density observed in the differences of eGCX thickness between IgE- and IgG1 PSA mice (**Figure 4C-D**). The structural analysis only showed changes in eGCX thickness of Sdx-treated mice (**Figure 4D**). Similarly to the TEM analysis, WGA intensity results significantly decreased in the endothelium and medial layer of both IgE-PSA and IgG1-PSA mice, but endothelial WGA intensity was only recovered in Sdx-treated IgE-PSA with no accompanying change to the medial layer (**Figure 4E**). Similarly to ASA mice, ESM-1 plasma levels were elevated in both IgE-PSA and IgG1-PSA mice without statistical modifications in groups treated with Sdx but stablishing a positive correlation with the drop in temperature (**Figure 4F-G**).

### eGCX shedding in an *in vitro* human endothelial/anaphylactic sera patients system

To translate the relevance of eGCX to humans, an *in vitro* human endothelial cells system was stablished. TEM analysis exhibited a dense layer of eGCX in HAECs that was significant reduced when HAECs were exposed to anaphylactic mediators (**Figure 5A-B**). Next, to investigate ESM-1 shedding from the eGCX, HMVEC-d were incubated with acute- and baseline-phase serum samples from patients with anaphylaxis. Following 2 h of exposure, the secretome of HMVEC-d treated with acute-phase serum showed a substantial increase in ESM-1 protein levels, providing compelling evidence that sera from anaphylactic patients induce ESM-1 release (**Figure 5C**). Next, to evaluate the potential of ESM-1 as a biomarker in anaphylaxis, circulating ESM-1 levels were measured in paired acute and baseline serum samples from anaphylactic patients and in serum from control subjects. The clinical characteristics of the patient cohort are summarised in **Table 1** and **Supplementary Table 1**. Significant differences in ESM-1 levels were observed between acute sera samples from anaphylactic patients compared to non-allergic controls (**Figure 5D**). Finally, to strengthen the potential of other eGCX components released into blood as human biomarkers, HA was measured. The analysis detected significant higher levels of HA in anaphylactic patients compared with those of controls (**Figure 5 E**).

**Figure 5.**
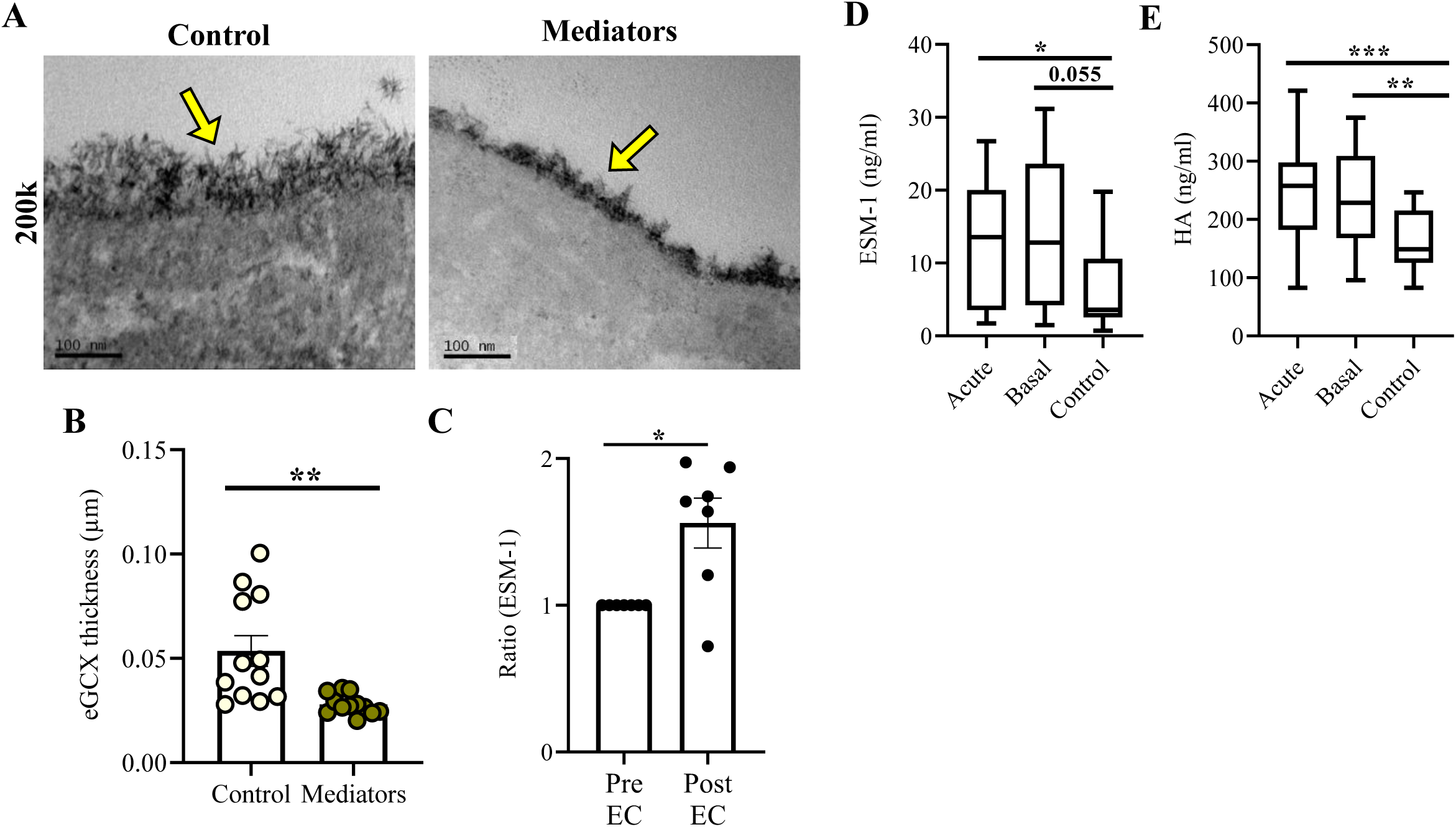
Endothelial glycocalyx components ESM-1 and HA are altered in human anaphylaxis. **A.** Representative TEM images of eGCX in HAEC cultures at 200k magnification. Yellow arrows point to eGCX**. B.** Quantification of eGCX thickness (µm) in control (n=2) and mediators (n=2). **C**. Increased ratio of ESM-1 levels in the secretome of serum from n=7 anaphylactic patients relative to their baseline levels after endothelial cell contact. ESM-1 (**D)** and (HA) (**E**) raw levels (ng/ml) in sera from 22 anaphylactic patients (acute and baseline) and 21 control subjects. eGCX: endothelial glycocalyx, TEM: transmission electronic microscopy, ESM-1: endothelial specific molecule 1, HA: hyaluronic acid, EC: endothelial cells. (*=*P*<0.05, **=*P*<0.01, ***=*P*<0.001).

**Table 1.** Characteristics of patients and their anaphylactic reactions. SEM: standard error of the mean; H1R: histamine receptor 1; H2R: histamine receptor 2.

|  | **Patients** |
| --- | --- |
| **Number** | 22 |
| **Sex (Female)** | 82% |
| **Age (years) ± SEM** | 43.09±2.65 |
| **Severity** |  |
| 2 (Moderate) | 59% |
| 3 (Severe) | 41% |
| **Symptoms** |  |
| Skin | 100% |
| Mucosa | 41% |
| Digestive | 50% |
| Respiratory | 86% |
| Nervous system | 27% |
| Cardiovascular | 41% |
| **Treatment** |  |
| Epinephrine | 73% |
| H1R Antagonist | 77% |
| H2R Antagonist | 9% |
| Corticosteroids | 86% |
| β2-adrenergic agonists | 5% |

## DISCUSSION

This study provides evidence that anaphylaxis triggers eGCX shedding, as demonstrated by *in vivo* animal studies showing decreased thickness of the aortic eGCX measured by TEM. This decrease correlated with the release of ESM-1 in the aorta and subsequent increase in plasma samples of ASA mice. eGCX shedding in IgE-PSA and IgG1-PSA aortas support the involvement of distinct immunological pathways inducing endothelial injury. Moreover, *in vitro* experiments showed that ESM-1 release is a direct consequence of eGCX shedding, as the secretome of *in vitro* human ECs exhibited significantly increased ESM-1 levels after contact with sera samples from anaphylactic patients. Changes in ESM-1 were also observed between acute sera samples from patients who had experienced anaphylaxis, and non-allergic subjects. Sdx prophylaxis reduced the severity of anaphylaxis in ASA and IgE-PSA mice and increased the survival rate of ASA, which was associated with a decrease in ESM-1 release into the bloodstream. Any prophylactic effect was seen in IgG1-PSA mice. This is the first study to describe the role played by eGCX in anaphylaxis pathophysiology, underscoring the potential of this structure as a mediator source and therapeutic target.

The eGCX is the primary endovascular barrier responsible for endothelial activation as demonstrated by increased *selp* and *vcam1* RNAs in ASA aorta mice. This hallmark is a well-established feature of the pathophysiology of most cardiovascular diseases, including atherosclerosis and acute inflammatory conditions such as sepsis.(15,42) In anaphylaxis, luminal eGCX degradation might be the starting point of endothelial dysfunction, triggering vascular permeability, loss of vascular tone, and heightened susceptibility to platelet adhesion, which contribute to the activation of coagulation systems, among others. A seminal study examining the structure of the eGCX in the myocardium of dogs in response to hypersensitivity reactions concluded that cytotoxic and immune complex-mediated reactions produced the most significant changes.(43) Our study provides a visualisation and quantification of the rupture of the fragile eGCX structure *in vivo* only 10 min after the challenge. Successful visualisation was achieved using Alcian blue (as a proteoglycan stabiliser) and lanthanum ions during glutaraldehyde fixation, instead of the more commonly used ruthenium red.(44) Thus, at the level of the thoracic aorta (approximately 60nm of thickness), the combination of TEM analysis and WGA staining showed eGCX shedding and proteoglycan degradation in ASA and both IgE-PSA and IgG1 PSA mice. In anaphylaxis, the high protease activity of some mediators degranulated from mast cells might proteolyze the eGCX architecture.(3) However, the oxidative environment created by NO or other mediators released by neutrophils or monocytes might also contribute to the fragmentation of the eGCX,(45) the loss of its components, and the endothelial dysfunction associated with anaphylaxis. Whether this occurs to a greater or lesser extent in all vascular beds and the number of molecular components released from the vascular wall remains unresolved. In different pathological settings, prevention of eGCX shedding appears beneficial.(46) (47) Restoration of a damaged eGCX is a therapeutic goal, primarily involving the administration of colloid and crystalloid solutions.(48) In general, fluid therapy is adopted in clinical interventions (49) instead of a specific treatment for eGCX protection or reconstruction. Clinical guidelines recommend fluid resuscitation for treatment of anaphylaxis, including refractory reactions.(7,8) However, Sdx, with its anti-inflammatory and anticoagulatory properties, protects the eGCX in sepsis.(19) Here, we demonstrate that Sdx improves survival in mouse models of anaphylaxis by mitigating reaction onset, including temperature drop and reaction severity without changes in hematocrit. In a previous study, 82% of ASA mice died within 60 to 90 min of i.v OVA challenge,(50) resembling the ASA model used in our study. However, we report that Sdx prophylaxis significantly improved survival and reaction severity in ASA. In addition, the selective protective effect of Sdx decreasing drop temperature in IgE-PSA mice points to a mechanism specific to IgE-mediated responses, suggesting that Sdx prophylaxis could mitigated IgE-mediated eGCX shedding, making it an attractive therapeutic tool.

RNA transcriptional activity in the aortic tissue of ASA mice revealed differential expression of genes related to eGCX components. *Synd1* exhibited a tendency to increase but was not regulated at this time point, although their potential contribution at later stages cannot be ruled out. However, the quick transcriptional upregulation of *esm-1* in the aortas of ASA mice appears to be driven by action of the amount of different pro-inflammatory factors and may even depend on anaphylaxis reaction time. Previous research has shown that *esm-1* mRNA increases in response to various pro-inflammatory cytokines and growth factors. Moreover, *esm-1* mRNA appears significantly elevated in response to hypoxia-inducible factor-1 α (HIF-1α).(51) Conversely, *esm-1* mRNA expression is down-regulated by interleukin-4 (IL-4), interferon-γ (IFN-γ), and phosphatidylinositol 3-kinase (PI3K).(52) At the protein level, ESM-1 staining was reduced in both the intimal and, even more, in the medial layers of ASA aortas compared with control samples. These findings suggest that ESM-1 is not only an eGCX component but also contributes to extracellular matrix (EM) organisation. The marked ESM-1 reduction observed here is consistent with previous *in vitro* studies demonstrating that acute-phase serum samples from anaphylactic patients decreases EM protein levels in human ECs.(53)

There is little doubt that ESM-1 plays a role in vascular cell biology and cardiovascular diseases, where elevated tissue expression or serum levels reflect endothelial activation.(30,31,51,54) Beyond the inflammatory properties in which ESM-1 is involved, an inverse correlation has been observed between serum ESM-1 levels and lipoprotein particles in Type 2 diabetes.(55) Serum samples from patients with severe anaphylaxis show decreased levels of HDL-C and LDL-C,(56) suggesting a possible inverse association between lipoproteins and ESM-1. In ASA and PSA mice, plasma ESM-1 levels are substantially increased, likely due to rapid shedding rather than the induction of vascular synthesis, as supported by *in vitro* studies showing ESM-1 in the secretome of ECs as a direct consequence of contact between these cells and human anaphylactic sera. Accordingly, differences in ESM-1 were observed in acute-phase sera samples from patients compared to non-allergic subjects. Although no differences were seen with their own baseline samples collected 14 days after the anaphylactic reaction. Certain limitations should be considered when interpreting these results. First, the small number of patient samples analysed restricts evaluation of possible treatment-induced adverse effects. These may alter ESM-1 levels, as most patients received medication prior to sample collection. Corticosteroids, other immunosuppressor drugs, statins, and hormones affect ESM-1 levels.(57,58) Second, sample timing varied between study systems, and mice sera samples were collected 10-20 min after challenge, while timing would varied in the case of patients. In addition, the *in vitro* human studies showed that anaphylactic mediators triggers eGCX disruption and ESM-1 accumulation within 2 h. Overall, eGCX shedding intensity likely varied depending on a plethora of intrinsic or extrinsic factors. Therefore, levels of other eGCX component, as HA, were evaluated and showed higher contents in anaphylactic samples than in controls. In relation, only one previous study showed an increase of hyaluronan in dog sera with suspected anaphylaxis due to insect envenomation.(59) As paired anaphylactic samples did not exhibit any changes in ESM-1 neither HA levels but were higher in both phases compared to non-allergic control samples, we cannot rule out ESM-1 and HA continuous release in the anaphylactic patient, suggesting that eGCX shedding in humans is not only a risk by exposing the endothelium but also contributes to chronic endothelial dysfunction and sustained vessel inflammation. Interestingly, our results show that Sdx prophylaxis specifically reduces ESM-1 levels in serum during ASA and IgE-PSA but not in IgG_1_-PSA. Although a positive correlation was found between drop temperature and ESM-1 release, Sdx treatment did not prevents ESM-1 release to the bloodstream neither severity. Overall, Sdx exhibit potential as a treatment to mitigate the paracrine effects of ESM-1.

In summary, our results indicate that eGCX shedding, particularly of ESM-1, plays a role in ASA and PSA. The stabilization of the eGCX with Sdx prophylaxis helps to reduce symptom severity in ASA and IgE-PSA and improves survival rates in these animals with a reduction in serum ESM-1 levels. These findings suggest that therapies based on GAGs and proteoglycans could mitigate these severe, life-threatening reactions.

## Abbreviations used

ASA: active systemic anaphylaxis
EC: endothelial cell
eGCX: endothelial glycocalyx
ESM-1: endothelial cell specific molecule 1(endocan)
GAG: glycosaminoglycan
HA: hyaluronic acid
IgE: immunoglobulin E
IgG1: immunoglobulin G1
PSA: passive systemic anaphylaxis
Sdx: Sulodexide
TEM: transmission electron microscopy
WGA: wheat germ agglutin

## Author Contribution Statement

LPG and SFB performed the experimental work with participation from IMC, ISJ, MJF-G, OL-P, SSA, IMC, and ENB. JJL and MV recruited human serum samples. LPG, SFB, and LMBC participated in the statistical analysis. VE and NMB coordinated the work, designed the experiments, and interpreted the results with participation from SFB and LPG. VE and NMB drafted the manuscript. All authors reviewed and approved the submitted manuscript.

### Acknowledgements

We are grateful to Maria Luisa Garcia Gil and Miriam Gonzalez Garcia from the National Center of electronic microscopy (Complutense University, Madrid) for technical assistance.

## References

1. Cardona V, Ansotegui IJ, Ebisawa M, El-Gamal Y, Rivas MF, Fineman S, et al. [World Allergy Organization Anaphylaxis Guidance 2020]. Arerugi. 2021;70(9):1211–34.

2. Nuñez-Borque E, Betancor D, Pastor-Vargas C, Fernández-Bravo S, Martin-Blazquez A, Casado-Navarro N, et al. Personalized diagnostic approach and indirect quantification of extravasation in human anaphylaxis. Allergy. 2023 Jan;78(1):202–13.

3. Nuñez-Borque E, Fernandez-Bravo S, Yuste-Montalvo A, Esteban V. Pathophysiological, Cellular, and Molecular Events of the Vascular System in Anaphylaxis. Front Immunol. 2022;13:836222.

4. Witjas FMR, van den Berg BM, van den Berg CW, Engelse MA, Rabelink TJ. Concise Review: The Endothelial Cell Extracellular Matrix Regulates Tissue Homeostasis and Repair. Stem Cells Transl Med. 2019 Apr;8(4):375–82.

5. Fernandez-Bravo S, Palacio Garcia L, Requena-Robledo N, Yuste-Montalvo A, Nuñez-Borque E, Esteban V. Anaphylaxis: Mediators, Biomarkers, and Microenvironments. J Investig Allergol Clin Immunol. 2022 Dec 15;32(6):419–39.

6. Claesson-Welsh L, Dejana E, McDonald DM. Permeability of the Endothelial Barrier: Identifying and Reconciling Controversies. Trends Mol Med. 2021 Apr;27(4):314–31.

7. Pouessel G, Dribin TE, Tacquard C, Tanno LK, Cardona V, Worm M, et al. Management of Refractory Anaphylaxis: An Overview of Current Guidelines. Clin Exp Allergy. 2024 July;54(7):470–88.

8. Muraro A, Worm M, Alviani C, Cardona V, DunnGalvin A, Garvey LH, et al. EAACI guidelines: Anaphylaxis (2021 update). Allergy. 2022 Feb;77(2):357–77.

9. Jedlicka J, Becker BF, Chappell D. Endothelial Glycocalyx. Crit Care Clin. 2020 Apr;36(2):217–32.

10. Radeva MY, Waschke J. Mind the gap: mechanisms regulating the endothelial barrier. Acta Physiol (Oxf). 2018 Jan;222(1).

11. Sieve I, Münster-Kühnel AK, Hilfiker-Kleiner D. Regulation and function of endothelial glycocalyx layer in vascular diseases. Vascul Pharmacol. 2018 Jan;100:26–33.

12. Nelson A, Berkestedt I, Schmidtchen A, Ljunggren L, Bodelsson M. Increased levels of glycosaminoglycans during septic shock: relation to mortality and the antibacterial actions of plasma. Shock. 2008 Dec;30(6):623–7.

13. Suzuki K, Okada H, Takemura G, Takada C, Kuroda A, Yano H, et al. Neutrophil Elastase Damages the Pulmonary Endothelial Glycocalyx in Lipopolysaccharide-Induced Experimental Endotoxemia. Am J Pathol. 2019 Aug;189(8):1526–35.

14. Sallisalmi M, Tenhunen J, Yang R, Oksala N, Pettilä V. Vascular adhesion protein-1 and syndecan-1 in septic shock. Acta Anaesthesiol Scand. 2012 Mar;56(3):316–22.

15. Uchimido R, Schmidt EP, Shapiro NI. The glycocalyx: a novel diagnostic and therapeutic target in sepsis. Crit Care. 2019 Jan 17;23(1):16.

16. Kataoka H, Ushiyama A, Akimoto Y, Matsubara S, Kawakami H, Iijima T. Structural Behavior of the Endothelial Glycocalyx Is Associated With Pathophysiologic Status in Septic Mice: An Integrated Approach to Analyzing the Behavior and Function of the Glycocalyx Using Both Electron and Fluorescence Intravital Microscopy. Anesth Analg. 2017 Sept;125(3):874–83.

17. Villalba N, Baby S, Yuan SY. The Endothelial Glycocalyx as a Double-Edged Sword in Microvascular Homeostasis and Pathogenesis. Front Cell Dev Biol. 2021;9:711003.

18. Andreozzi GM. Sulodexide in the treatment of chronic venous disease. Am J Cardiovasc Drugs. 2012 Apr 1;12(2):73–81.

19. Ying J, Zhang C, Wang Y, Liu T, Yu Z, Wang K, et al. Sulodexide improves vascular permeability via glycocalyx remodelling in endothelial cells during sepsis. Front Immunol. 2023;14:1172892.

20. Callas DD, Hoppensteadt DA, Jeske W, Iqbal O, Bacher P, Ahsan A, et al. Comparative pharmacologic profile of a glycosaminoglycan mixture, Sulodexide, and a chemically modified heparin derivative, Suleparoide. Semin Thromb Hemost. 1993;19 Suppl 1:49–57.

21. Carroll BJ, Piazza G, Goldhaber SZ. Sulodexide in venous disease. J Thromb Haemost. 2019 Jan;17(1):31–8.

22. Mannello F, Ligi D, Raffetto JD. Glycosaminoglycan sulodexide modulates inflammatory pathways in chronic venous disease. Int Angiol. 2014 June;33(3):236–42.

23. Hoppensteadt DA, Fareed J. Pharmacological profile of sulodexide. Int Angiol. 2014 June;33(3):229–35.

24. Lassalle P, Molet S, Janin A, Heyden JV, Tavernier J, Fiers W, et al. ESM-1 is a novel human endothelial cell-specific molecule expressed in lung and regulated by cytokines. J Biol Chem. 1996 Aug 23;271(34):20458–64.

25. Bechard D, Meignin V, Scherpereel A, Oudin S, Kervoaze G, Bertheau P, et al. Characterization of the secreted form of endothelial-cell-specific molecule 1 by specific monoclonal antibodies. J Vasc Res. 2000;37(5):417–25.

26. Grigoriu BD, Depontieu F, Scherpereel A, Gourcerol D, Devos P, Ouatas T, et al. Endocan expression and relationship with survival in human non-small cell lung cancer. Clin Cancer Res. 2006 Aug 1;12(15):4575–82.

27. Kirwan RP, Leonard MO, Murphy M, Clark AF, O’Brien CJ. Transforming growth factor-beta-regulated gene transcription and protein expression in human GFAP-negative lamina cribrosa cells. Glia. 2005 Dec;52(4):309–24.

28. Abid MR, Yi X, Yano K, Shih SC, Aird WC. Vascular endocan is preferentially expressed in tumor endothelium. Microvasc Res. 2006 Nov;72(3):136–45.

29. Balta S, Balta I, Mikhailidis DP. Endocan: a new marker of endothelial function. Curr Opin Cardiol. 2021 July 1;36(4):462–8.

30. Balta S, Mikhailidis DP, Demirkol S, Ozturk C, Celik T, Iyisoy A. Endocan: A novel inflammatory indicator in cardiovascular disease? Atherosclerosis. 2015 Nov;243(1):339–43.

31. Kali A, Shetty KSR. Endocan: a novel circulating proteoglycan. Indian J Pharmacol. 2014;46(6):579–83.

32. Klisic A, Kotur-Stevuljevic J, Gluscevic S, Sahin SB, Mercantepe F. Biochemical markers and carotid intima-media thickness in relation to cardiovascular risk in young women. Sci Rep. 2024 Oct 21;14(1):24776.

33. De Freitas Caires N, Gaudet A, Portier L, Tsicopoulos A, Mathieu D, Lassalle P. Endocan, sepsis, pneumonia, and acute respiratory distress syndrome. Crit Care. 2018 Oct 26;22(1):280.

34. Bessa J, Albino-Teixeira A, Reina-Couto M, Sousa T. Endocan: A novel biomarker for risk stratification, prognosis and therapeutic monitoring in human cardiovascular and renal diseases. Clin Chim Acta. 2020 Oct;509:310–35.

35. Daiki K, Kanada Y, Nagata A, Taruno K, Igarashi K, Yamochi T, et al. Blood endocan as a biomarker for breast cancer recurrence. Cancer Biomark. 2024;41(2):145–54.

36. Behnoush AH, Khalaji A, Ghasemi H, Tabatabaei GA, Samavarchitehrani A, Vaziri Z, et al. Endocan as a biomarker for acute respiratory distress syndrome: A systematic review and meta-analysis. Health Sci Rep. 2024 Sept;7(9):e70044.

37. Scherpereel A, Depontieu F, Grigoriu B, Cavestri B, Tsicopoulos A, Gentina T, et al. Endocan, a new endothelial marker in human sepsis. Crit Care Med. 2006 Feb;34(2):532–7.

38. Sarrazin S, Adam E, Lyon M, Depontieu F, Motte V, Landolfi C, et al. Endocan or endothelial cell specific molecule-1 (ESM-1): a potential novel endothelial cell marker and a new target for cancer therapy. Biochim Biophys Acta. 2006 Jan;1765(1):25–37.

39. Ballesteros-Martinez C, Mendez-Barbero N, Montalvo-Yuste A, Jensen BM, Gomez-Cardenosa A, Klitfod L, et al. Endothelial Regulator of Calcineurin 1 Promotes Barrier Integrity and Modulates Histamine-Induced Barrier Dysfunction in Anaphylaxis. Front Immunol. 2017;8:1323.

40. Sampson HA, Muñoz-Furlong A, Campbell RL, Adkinson NF, Bock SA, Branum A, et al. Second symposium on the definition and management of anaphylaxis: summary report--Second National Institute of Allergy and Infectious Disease/Food Allergy and Anaphylaxis Network symposium. J Allergy Clin Immunol. 2006 Feb;117(2):391–7.

41. Brown SGA. Clinical features and severity grading of anaphylaxis. J Allergy Clin Immunol. 2004 Aug;114(2):371–6.

42. Ahn SJ, Le Master E, Granados ST, Levitan I. Impairment of endothelial glycocalyx in atherosclerosis and obesity. Curr Top Membr. 2023;91:1–19.

43. Popovich LF, Sagach VF, Moibenko AA. Comparative study of morphological changes in the myocardium after different types of allergic reaction in coronary vessels. Exp Pathol. 1988;33(2):109–17.

44. Mukai S, Takaki T, Nagumo T, Sano M, Kang D, Takimoto M, et al. Three-dimensional electron microscopy for endothelial glycocalyx observation using Alcian blue with silver enhancement. Med Mol Morphol. 2021 June;54(2):95– 107.

45. Milusev A, Rieben R, Sorvillo N. The Endothelial Glycocalyx: A Possible Therapeutic Target in Cardiovascular Disorders. Front Cardiovasc Med. 2022;9:897087.

46. Cerny V, Astapenko D, Brettner F, Benes J, Hyspler R, Lehmann C, et al. Targeting the endothelial glycocalyx in acute critical illness as a challenge for clinical and laboratory medicine. Crit Rev Clin Lab Sci. 2017 Aug;54(5):343–57.

47. Coccheri S, Mannello F. Development and use of sulodexide in vascular diseases: implications for treatment. Drug Des Devel Ther. 2013 Dec 24;8:49–65.

48. Becker BF, Chappell D, Bruegger D, Annecke T, Jacob M. Therapeutic strategies targeting the endothelial glycocalyx: acute deficits, but great potential. Cardiovasc Res. 2010 July 15;87(2):300–10.

49. Myburgh JA, Mythen MG. Resuscitation fluids. N Engl J Med. 2013 Sept 26;369(13):1243–51.

50. Cauwels A, Janssen B, Buys E, Sips P, Brouckaert P. Anaphylactic shock depends on PI3K and eNOS-derived NO. J Clin Invest. 2006 Aug;116(8):2244–51.

51. Sun H, Zhang H, Li K, Wu H, Zhan X, Fang F, et al. ESM-1 promotes adhesion between monocytes and endothelial cells under intermittent hypoxia. J Cell Physiol. 2019 Feb;234(2):1512–21.

52. Rennel E, Mellberg S, Dimberg A, Petersson L, Botling J, Ameur A, et al. Endocan is a VEGF-A and PI3K regulated gene with increased expression in human renal cancer. Exp Cell Res. 2007 Apr 15;313(7):1285–94.

53. Yuste-Montalvo A, Fernandez-Bravo S, Oliva T, Pastor-Vargas C, Betancor D, Goikoetxea MJ, et al. Proteomic and Biological Analysis of an In Vitro Human Endothelial System in Response to Drug Anaphylaxis. Front Immunol. 2021;12:692569.

54. Kumar SK, Mani KP. Proinflammatory signaling mechanism of endocan in macrophages: Involvement of TLR2 mediated MAPK-NFkB pathways. Cytokine. 2024 Mar;175:156482.

55. Klisic A, Kavaric N, Vujcic S, Mihajlovic M, Zeljkovic A, Ivanisevic J, et al. Inverse association between serum endocan levels and small LDL and HDL particles in patients with type 2 diabetes mellitus. Eur Rev Med Pharmacol Sci. 2020 Aug;24(15):8127–35.

56. Fernandez-Bravo S, Canyelles M, Martín-Blázquez A, Borràs C, Nuñez-Borque E, Palacio-García L, et al. Impaired high-density lipoprotein function and endothelial barrier stability in severe anaphylaxis. J Allergy Clin Immunol. 2024 Sept;154(3):827–32.

57. Celýk T, Balta S, Karaman M, Ahmet Ay S, Demırkol S, Ozturk C, et al. Endocan, a novel marker of endothelial dysfunction in patients with essential hypertension: comparative effects of amlodipine and valsartan. Blood Press. 2015 Feb;24(1):55–60.

58. Potje SR, Martins NS, Benatti MN, Rodrigues D, Bonato VLD, Tostes RC. The effects of female sexual hormones on the endothelial glycocalyx. Curr Top Membr. 2023;91:89–137.

59. Turner K, Boyd C, Rossi G, Sharp CR, Claus MA, Francis A, et al. Allergy, inflammation, hepatopathy and coagulation biomarkers in dogs with suspected anaphylaxis due to insect envenomation. Front Vet Sci. 2022;9:875339.

