## Supplemental material for "Attenuation of endothelial glycocalyx shedding and endocan modulation by Sulodexide in murine models of anaphylaxis"

### SUPPLEMENTARY MATERIAL

### Methods

#### Systemic anaphylaxis murine models

Active systemic anaphylaxis (ASA) and passive systemic anaphylaxis (PSA) models were carried out in C57BL/6 mice in accordance with guidelines and recommendations. For ASA, mice were sensitized with an intraperitoneal injection of 1mg BSA (Sigma-Aldrich) and 300ng Pertussis Toxin (Merck) as adjuvant in saline solution. After 14 days, mice were challenged with an intravenous (i.v.) injection of 2mg BSA in saline solution. On the other hand, IgE- and IgG-dependent PSA were carried out. Mice were sensitized with 20ug IgE anti-DNP monoclonal antibodies (mAb) (Merck) or 100ug IgG1 anti-DNP mAb (Across Biosystems). After 24h, mice were challenged with an i.v. injection of 1mg DNP-HSA (Human serum albumin) (Merck).

#### Preparation of samples for electron microscopy

For murine aortic samples preparation, after saline solution perfusion, animals were fixed with a fixative solution containing 2.5% paraformaldehyde (PFA; Electron Microscopy Science), 2.5% glutaraldehyde (Electron Microscopy Science), and 30 mM magnesium chloride ($MgCl_2$; Sigma). Perfusion was completed with a solution consisting of the previous one plus 0.05% Alcian Blue (AB; Sigma), which provides specific staining for acidic polysaccharides such as glycosaminoglycans (GAGs). After perfusion, the aortas were isolated and divided into 3–4-millimetre fragments, which were immersed in the fixative solution containing 0.05% AB at 4ºC overnight (ON). On the other hand, for cell culture sample preparation, after *in vitro* assays, cells were fixed with the fixative solution for 10 minutes (min) at room temperature (RT). Then, they were fixed for another 10min with the fixative solution containing 0.05% AB, and finally they were also immersed ON with this last solution. The following day, both the aortic fragments or cell samples were washed in saline solution and post-fixed with 1% osmium tetroxide (Electron Microscopy Science) and 1% lanthanum tetroxide (Sigma Aldrich) for 1 hour (h) at RT. After washing again in saline solution, the samples were dehydrated in increasing concentrations of ethanol (30%, 50%, 70%, 80%, 90%, 95%, 100%) for 15min at RT each. Then, tissues and cultures were gradually embedded in Spurr resin by immersing them in solutions with increasing resin concentration in 100% ethanol (3:1, 1:1, 1:1, 1:3, and pure resin) for 1h at RT each. Samples were left in pure resin at RT ON and slides were observed using electron microscopy. Images were taken in the TEM Department of the National center of electronic microscopy (Complutense University, Madrid).

**RT-qPCR**

All primers were purchased from Eurofins Genomics and designed according to the following sequences: *esm1* (forward primer (fw): 5' CTG GAG CGC CAA ATA TGC G 3'; reverse primer (rv): 5' TGA GAC TGT ACG GTA GCA GGT 3'), *synd1* (fw: 5'CCT TGT CAG GGT AGA CAG CCT 3'; rv: 5' GAC AGA GGT AAA AGC AGT CTC G 3'), *synd4* (fw: 5' CAT CTT TGA GAG AAC TGA GGT CTT 3'; rv: 5' CCT TCT TCT TCA TGC GGT ACA 3'), *acan* (fw: 5' CGC CAC TTT CAT GAC CGA GA 3'; rv: 5' TCA TTC AGA CCG ATC CAC TGG TAG 3'), *gpc1* (fw: 5' GGA GAG CGC ACT CCA TGA C 3'; rv: 5' CTC AGC ATA TAC GTC CCG GAA 3'), *hs2st1* (fw: 5' TAT GAT GCC GCC CAA GTT G 3'; rv: 5' CTG TTC AAT TTC TCG GAC TTC GT 3'), *ndst2* (fw: 5' CTG CTG ATT GGT TTC AGT CTT 3'; rv: 5' CCA CTG CTA CTA CAG TCT CCC 3'), *selp. (*fw*: 5´-GAA AGG GCT GAT TGT GAC CCC-3´;* rv*: 5´-AGT AGT TCC GCA CTG GGT ACA-3´.* 18S rRNA (fw: 5' CCG TCG TAG TTC CGA CCA TAA 3'; rv: 5' CAG CTT TGC AAC CAT ACT CCC 3') was used as an internal control. Reactions were incubated for 2 min at 50°C followed by 5 min at 95°C. Then they were run over 40 amplification cycles (95°C for 15 seconds, 60°C for 1 min and 72ºC for 40 seconds) followed by a dissociation stage (95ºC for 15 seconds, 60ºC for 20 seconds, and 98ºC for 15 seconds). The quantification *vcam1* (Mm01320970_m1) and 18s rRNA (4310893E) was performed with TaqMan Gene expression assays from Applied Biosystems. Reactions were incubated for 20 seconds at 95ºC and then run over 40 amplification cycles (95ºC for 3 seconds and 60ºC for 30 seconds).

**Human endothelial cell cultures**

Human dermal microvascular endothelial cells (HMVEC-d) and human aortic endothelial cells (HAEC) were acquired from Lonza (catalog number CC-2543 and K3CC-2535, respectively). Primary cell cultures were grown and expanded using EGM medium (EGM-2 MV Bullet Kit) supplemented with fetal bovine serum (FBS), hydrocortisone, human basic fibroblast growth factor (hFGF-B), vascular endothelial growth factor (VEGF), recombinant insulin-like growth factor-I analogue with the substitution of arginine for glutamine at position 3 (R3-IGF-1), ascorbic acid, human epidermal growth factor (hEGF), and gentamicin sulfate/amphotericin (GA-1000) (Lonza). The medium was further enriched with heparin and endothelial cell growth factor (ECGF).

### Supplementary figures legend

**Supplementary Figure 1. A**. Representative images of WGA (red), CD31 (white) and DAPI (blue) staining in aortic samples. **B**. WGA intensity profiles. **C.** Correlation between temperature decrease (degrees) and WGA intensity staining. **D**. Correlation between temperature decrease (degrees) and hematocrit percentage. Control (n=4) and ASA (n=8) mice. WGA: wheat germ agglutinin, ASA: active systemic anaphylaxis.

**Supplementary Figure 2**. Representative images of WGA (red), CD31 (green) and DAPI (blue) staining in aortic samples. WGA: wheat germ agglutinin, ASA: active systemic anaphylaxis, Sdx: sulodexide.

**Supplementary Figure 3. A.** Single heatmap for visualizing data for endothelial and eGCX genes. Numbers represent medium values of gene expression in aortas from control (n=4) and ASA (n=14) mice. Warm and cold colours denote high and low levels of gene expression, respectively. **B.** Different panels show the quantification of detected, *selp, vcam1*, *acan, synd1, synd4, gpc1, ndst2,* and *hs2st1* mRNA expression (*=$P$<0.05). Esm-1: endothelial specific molecule 1, selp: p-selectin; vcam1; vascular cell adhesion protein 1, acan: aggrecan, synd1: syndecan 1, synd4: syndecan 4, gpc1: glypican 1, hs2st: heparan sulfate 2-O-sulfotransferase 1, ndst2: N-deacetylase and N-sulfotransferase 2, ASA: active systemic anaphylaxis. **C.** Representative images of ESM-1 (green), and DAPI (blue) immunofluorescences in aortic samples.

**Supplementary Figure 4.** Representative images of WGA (red), CD31 (green) and DAPI (blue) staining in aortic samples. WGA: wheat germ agglutinin, Sdx: sulodexide.

### Supplementary Table

| Patient | | Signs and symptoms | | | | | | | | | | Tryptase | | Treatment | | | | |
|---|---|---|---|---|---|---|---|---|---|---|---|---|---|---|---|---|---|---|
| Sex | Age | Sk | Mu | Di | Re | Cv/Ne | SBP | DBP | Hr | $SatO_2$ | Severity | Acute | Basal | Ep | β2A | H1AR | H2AR | Cs |
| F | 33 | x | x | x | x | x | 120 | 95 | - | 95% | 2 (Moderate) | 7.29 | 3.51 | x | | x | | x |
| F | 37 | x | | x | x | | 130 | 85 | 90 | 99% | 2 (Moderate) | 5.39 | 5.4 | x | | x | | x |
| F | 65 | x | x | x | x | x | 123 | 72 | 68 | 95% | 2 (Moderate) | 6.94 | 5.37 | | | x | | x |
| F | 51 | x | | | | x | 120 | 80 | 90 | 95% | 2 (Moderate) | 24.9 | 9.33 | | | x | | x |
| F | 49 | x | x | x | x | | 130 | 80 | 70 | 94% | 2 (Moderate) | 5.32 | 5.82 | x | | | | x |
| M | 50 | x | x | x | x | x | 90 | 60 | 90 | 92% | 3 (Severe) | 12 | 8.63 | x | | x | | x |
| F | 25 | x | | | x | | 110 | 60 | 85 | - | 2 (Moderate) | 3.6 | 4.74 | x | x | x | | |
| M | 47 | x | | | | x | 170 | 80 | 78 | 89% | 3 (Severe) | 9.8 | 6.77 | x | | | x | x |
| F | 22 | x | | x | x | x | 120 | 75 | 130 | 91% | 3 (Severe) | 5.56 | 4.49 | x | | x | | x |
| F | 47 | x | | x | x | x | 125 | 65 | 86 | 97% | 3 (Severe) | 15.2 | 8.48 | x | | x | | x |
| F | 37 | x | | x | x | x | 150 | 90 | 100 | 94% | 2 (Moderate) | 10.8 | 7.35 | x | | | | |
| F | 59 | x | | x | | x | 50 | 30 | 127 | 100% | 3 (Severe) | 28.4 | 8.1 | x | | | | |
| M | 41 | x | x | | x | x | 142 | 82 | 114 | 100% | 2 (Moderate) | 8.01 | 4.48 | | | x | | x |
| F | 38 | x | x | x | x | | 71 | 45 | 130 | 92% | 3 (Severe) | 6.63 | 3.46 | x | | x | | x |
| F | 42 | x | | | x | | 105 | 75 | 76 | 100% | 2 (Moderate) | 7.98 | 5.73 | x | | x | | x |
| F | 44 | x | x | | x | | 105 | 66 | 68 | 99% | 2 (Moderate) | 4.8 | 4.08 | | | x | | x |
| F | 47 | x | | | x | | 127 | 88 | 80 | 100% | 2 (Moderate) | 4.52 | 5.01 | | | x | | x |
| F | 27 | x | x | | x | | 120 | 70 | 86 | 100% | 2 (Moderate) | 2,18 | 3,25 | x | | x | | x |
| M | 62 | x | x | | x | | 141 | 88 | 86 | 97% | 2 (Moderate) | 3,4 | 3,95 | x | | x | | x |
| F | 63 | x | | x | x | | 70 | 50 | | 90% | 3 (Severe) | 12,9 | 3,73 | x | | x | | x |
| F | 26 | x | x | | x | | 50 | 25 | | 89% | 3 (Severe) | 23,7 | 6,26 | x | | | | x |
| F | 36 | x | | | x | | 111 | 72 | 117 | 80% | 3 (Severe) | 38,6 | 4,17 | x | | x | x | x |

**Supplementary Table 1.** Clinical features of anaphylactic reactions included in this study. The study population included 22 anaphylactic patients ranging in age from 22 to 65 years. The criteria of severity details of each individual case (criteria of severity) were applied based on clinical symptoms and according to the grading system established in the classification by Brown (**1**) being 2 moderate and 3 severe. Severe anaphylaxis includes those with cardiovascular involvement and hypotension. Sex: M (male), F (female). Symptoms: Sk (skin), Mu (mucous), Di (digestive), Re (respiratory), Cv (cardiovascular), Ne (neurological). Signs: SBP (systolic blood pressure), DBP (diastolic blood pressure), Hr (heart rate), $SatO_2$ (oxygen saturation). Treatments: Ep (epinephrine), β2A (β2-adrenergic agonist), H1RA (histamine 1-receptor antagonist), H2RA (histamine 2-receptor antagonist), Cs (corticosteroids). (**1**) Brown SGA. Clinical features and severity grading of anaphylaxis. J Allergy Clin Immunol. 2004;114(2):371-376.

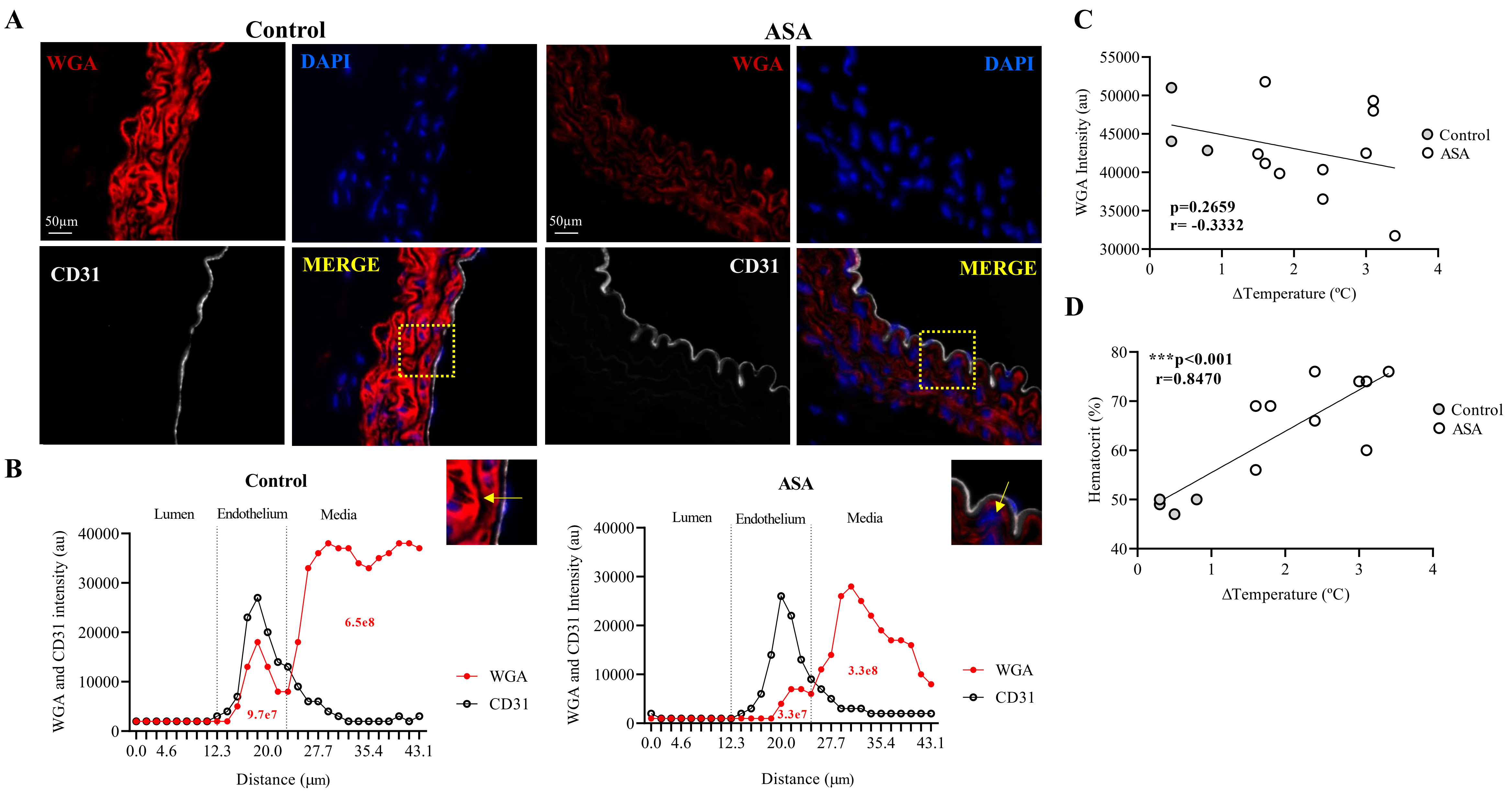


**Supplemental figure 1**

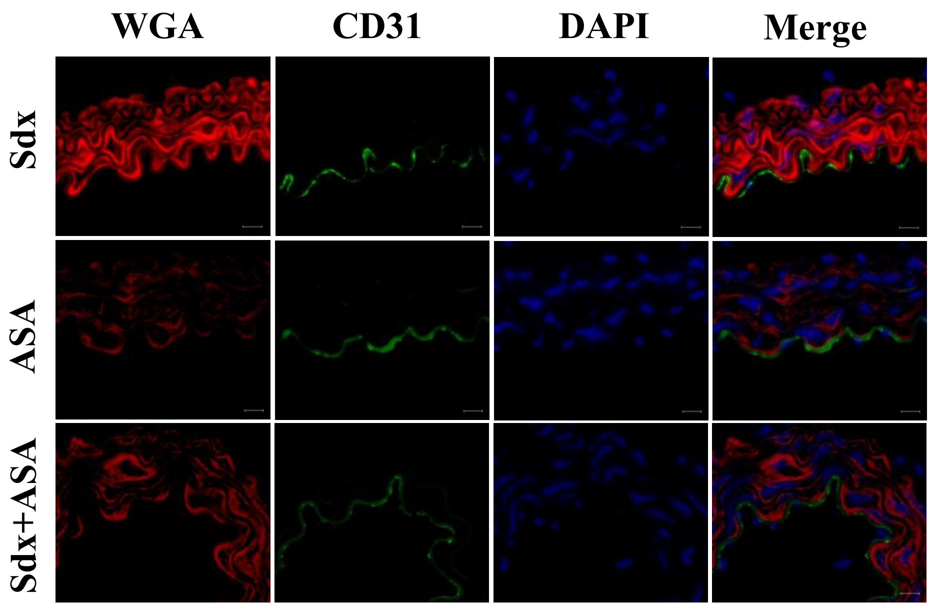


**Supplemental figure 2**

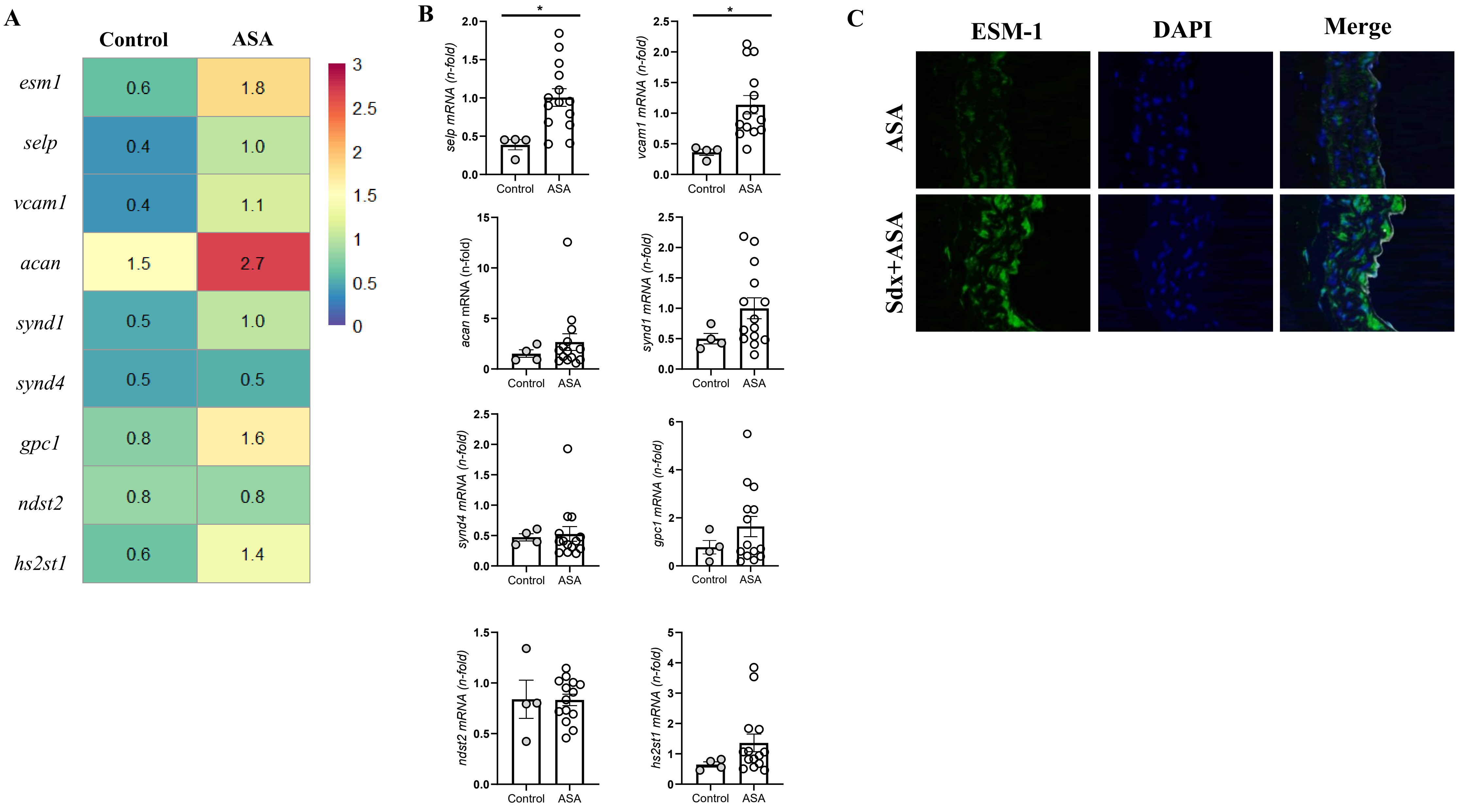


**Supplemental figure 3**

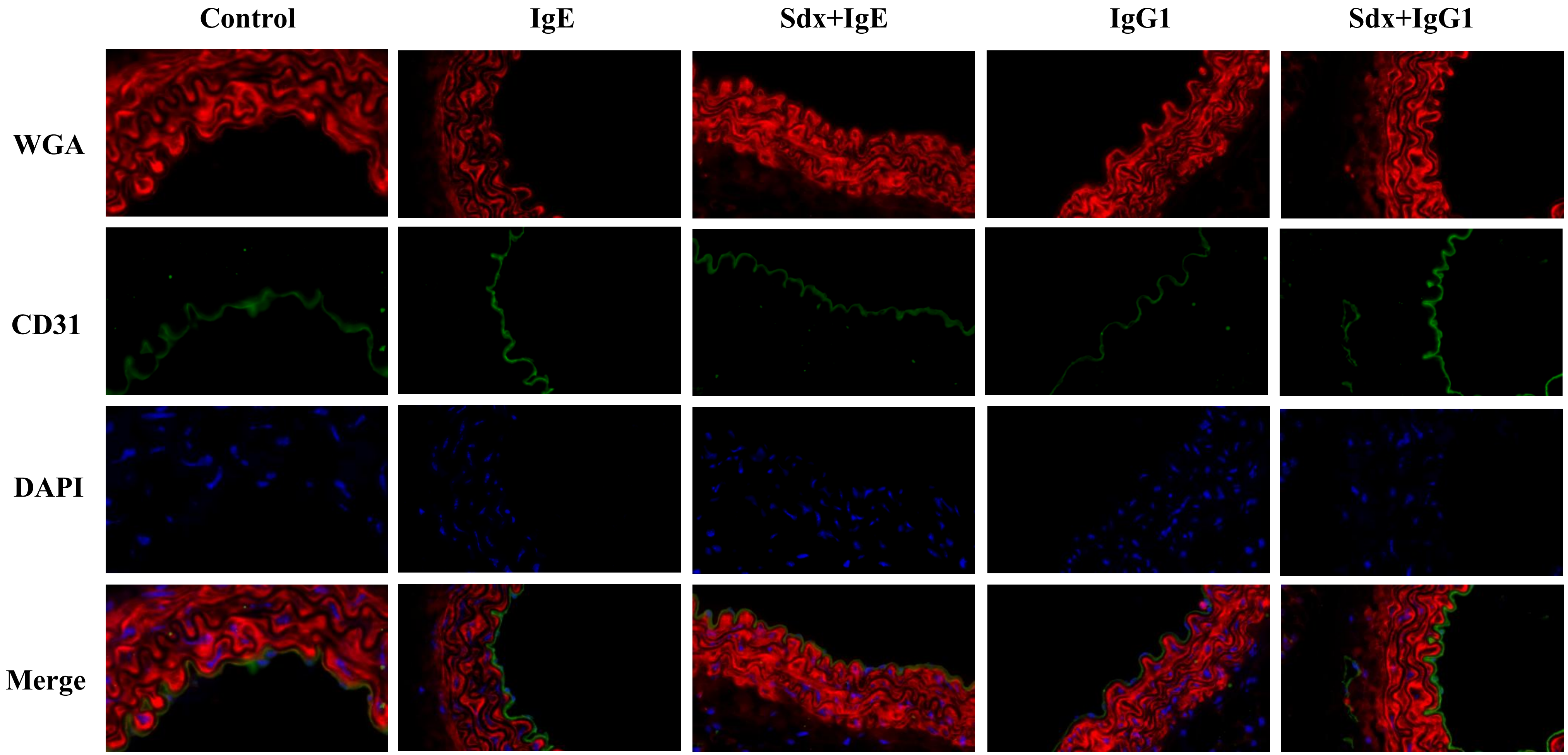


**Supplemental figure 4**
